# OVOL2 induces mesenchymal-to-epithelial transition in fibroblasts and enhances reprogramming to epithelial lineages

**DOI:** 10.1101/577692

**Authors:** Kazuhide Watanabe, Ye Liu, Shuhei Noguchi, Madeleine Murray, Jen-Chien Chang, Mami Kishima, Hajime Nishimura, Kosuke Hashimoto, Aki Minoda, Harukazu Suzuki

## Abstract

Mesenchymal-to-epithelial transition (MET) is an important step in cell reprogramming from fibroblasts (a cell type frequently used for this purpose) to various epithelial cell types. However, the mechanism underlying MET induction in fibroblasts remains to be understood. The present study aimed to identify the transcription factors (TFs) that efficiently induce MET in dermal fibroblasts. OVOL2 was identified as a potent inducer of key epithelial genes, and OVOL2 cooperatively enhanced MET induced by HNF1A, TP63, and KLF4, which are known reprogramming TFs to epithelial lineages. In TP63/KLF4-induced keratinocyte-like cell-state reprogramming, OVOL2 greatly facilitated the activation of epithelial and keratinocyte-specific genes. This was accompanied by enhanced changes in chromatin accessibility across the genome. Mechanistically, motif enrichment analysis revealed that the target loci of KLF4 and TP63 become accessible upon induction of TFs, whereas the OVOL2 target loci become inaccessible. This indicates that KLF4 and TP63 positively regulate keratinocyte-associated genes whereas OVOL2 suppresses fibroblast-associated genes. The exogenous expression of OVOL2 therefore disrupts fibroblast lineage identity and facilitates fibroblast cell reprogramming into epithelial lineages cooperatively with tissue-specific reprogramming factors. Identification of OVOL2 as a MET inducer and an epithelial reprogramming enhancer in fibroblasts provides new insights into cellular reprogramming improvement for future applications.

## INTRODUCTION

Fibroblasts are mesenchymal cells most frequently used for cell reprogramming. Several studies have established the importance of mesenchymal-to-epithelial transition (MET) to reprogram fibroblasts into various epithelial cell types. The initial stages of induced pluripotent stem cell (iPSC) reprogramming from fibroblasts require MET, and its efficiency is significantly suppressed by MET inhibition ^1–3^. MET is also involved in fibroblasts’ direct cell reprogramming into an epithelial hepatocyte-like state ^4^. In fact, the mesenchymal identity of fibroblasts is maintained/protected by multiple mesenchymal (M)–epithelial (E) barriers, including DNA methylation ^5^, barrier kinases ^6^, and microRNAs ^7^. This suggests that activating the intrinsic MET program may enhance the efficiency of cellular reprogramming of fibroblasts into epithelial cell lineages. However, MET induction in fibroblasts has not been fully explored.

Epithelial-to-mesenchymal transition (EMT), an opposite process of MET, is involved in developmental dynamic tissue morphogenesis, as well as in fibrosis, invasion, metastasis, stemness, and chemo-resistance of tumor cells under pathological conditions ^8^. Therefore, the underlying mechanism of EMT has been extensively studied over decades, and several transcription factors such as SNAIL/SLUG, ZEB, and TWIST families have been identified as EMT-inducing transcription factors (EMT-TFs) ^9^. On the other hand, MET has recently drawn attention as an attractive target for cancer therapy ^10^ and several TFs have been found to exhibit MET-inducing activity (MET-TFs). In various cancer cells, GRHL2 is associated with epithelial phenotype ^11^ and activates epithelial enhancer subsets during the early differentiation of pluripotent cells ^12^. The OVOL family induces MET in several cancer cell types ^13^ and protects the epithelial identity of normal epithelial cells ^14,15^, whereas the ELF family is reported to inhibit EMT in ovarian cancer ^16^ and mammary epithelial cells ^17^. GATA3 initiates MET in breast cancer cells as a pioneer TF ^18^, whereas FOXA1 and FOXA3 maintain the expression of key epithelial genes in colorectal cancer cells ^19^ and KLF4 exhibits MET-inducing activity during iPSC reprogramming ^20^. However, MET-TF(s) that can efficiently induce MET in fibroblasts remain to be identified.

The present study aimed to identify TFs that can activate MET in dermal fibroblasts. We identified OVOL2 as a potent MET-TF enhancer, which can cooperatively induce MET with known reprogramming TFs, HNF1A, TP63, and KLF4. We also demonstrated that OVOL2 facilitates KLF4/TP63-induced keratinocyte-like cell state in fibroblasts. The Assay for Transposase-Accessibility of Chromatin (ATAC)-seq analysis disclosed a mechanism where KLF4/TP63 and OVOL2 play different roles in the regulation of chromatin accessibility in their target loci.

## RESULTS

### Selection of MET-TF candidates in fibroblasts

To identify the TFs promoting MET in fibroblasts, we first selected candidate MET-TFs among 1995 TF genes annotated by functional annotation of mammalian genome 5 (FANTOM5) ^21^, based on a positive or negative expression correlation with E or M markers, respectively. *CDH1* and *VIM* were used as representative E and M marker genes, respectively, since the expression of these genes showed the strongest anti-correlation among typical E and M markers in two public gene expression databases: the Cancer Cell Line Encyclopedia (CCLE, https://portals.broadinstitute.org/ccle/) and the FANTOM5 (Figure S1). We calculated and plotted the Pearson correlation coefficient *r*, between the expression of TF genes and the expression of *CDH1* or *VIM*, across 1038 CCLE microarrays for cancer cell lines representing various degrees of E and M states (Figure 1A). As expected, known MET-TF genes, the *OVOL* ^13^ and *GRHL* ^11^ families, exhibited a strongly positive correlation with *CDH1* and a negative correlation with *VIM*, whereas known EMT-TF genes, the *SNAI, TWIST*, and *ZEB* families, exhibited opposite correlations, thus confirming the validity of this approach. Ten candidate MET-TFs were selected from this plot (i.e., ANKRD22, EHF, ELF3, FOXA1, GRHL1/2/3, IRF6, OVOL2, and ZNF165). In addition, GATA3 ^18^, FOXA3 ^19^, HNF1A ^22^, KLF4 ^20^, and TP63 ^23^ were selected on the basis of their previously reported involvement in MET activation or EMT suppression. A correlation between these candidate factors and the E state was also observed in 1829 FANTOM5 data for normal primary cell types (Figure S2), where YBX2 was selected on the basis of its strong association with the E state. This resulted in a list of 16 candidate MET-TFs (Table 1).

**Figure 1.**
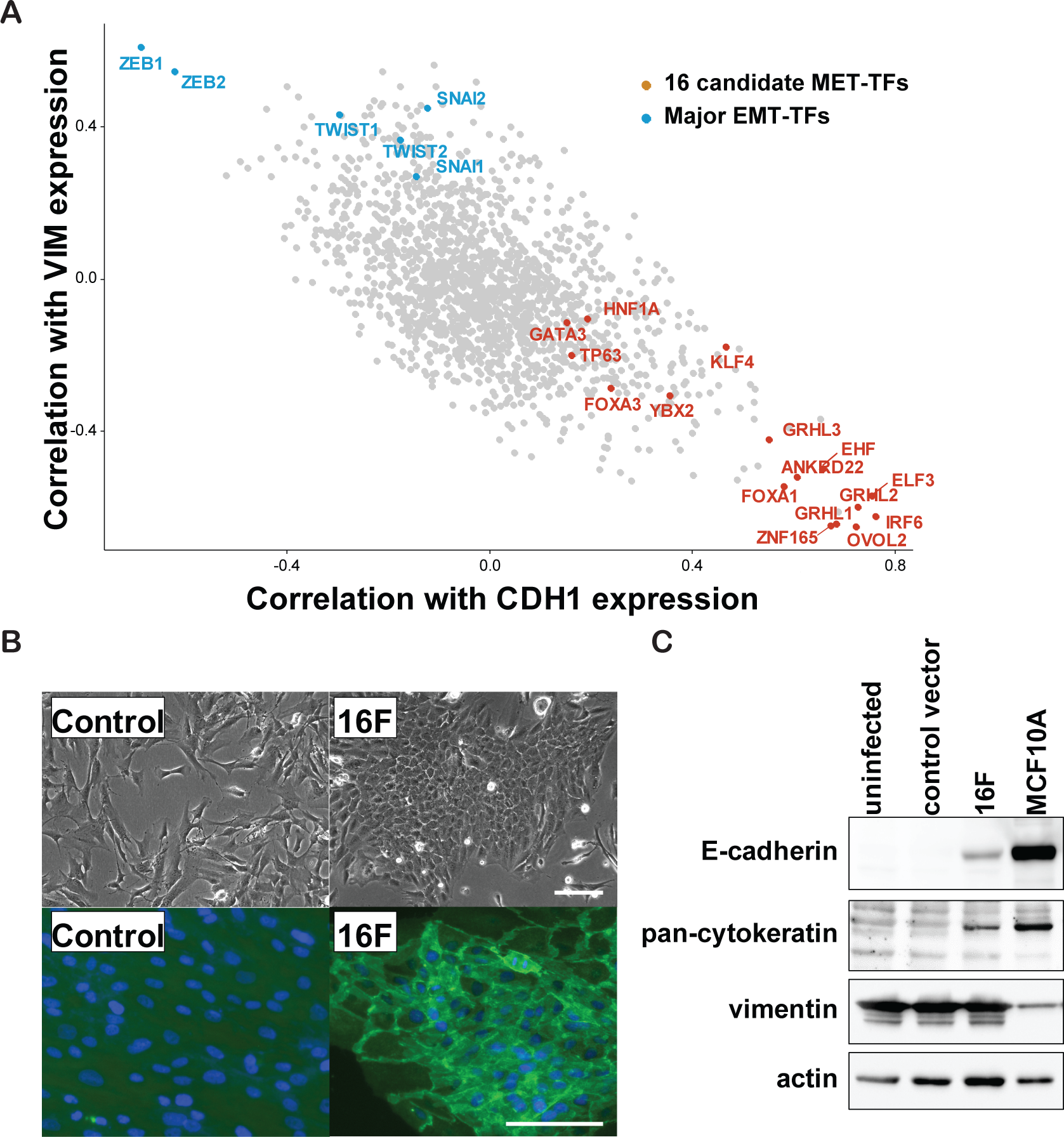
Screening of TF combinations to induce MET in fibroblasts. (**A**) Expression correlation analysis of FANTOM5-defined TFs in CCLE microarray dataset. Among 1995 FANTOM5-defined TFs, 1681 TF transcripts were identified from the CCLE datasets. The Pearson correlation coefficient *r* was calculated between each transcript and *CDH1* or *VIM* across 1038 CCLE microarrays of cancer cell lines. Representative EMT-TFs (blue) and 16 candidate MET-TFs (red) were labeled. (**B**) E-state induction by the 16 candidate MET-TFs. TFs were introduced by lentiviral transduction (20 MOI) in human neonatal dermal fibroblasts (HNDFs). The images were taken at post-transduction day 10. Upper panel: phase-contrast images. Lower panel: immunofluorescent staining of β-catenin and DAPI showing typical epithelial junction structures. Scale bar = 50 μm. (**C**) Detection of E markers (E-cadherin and pan-cytokeratin) and M markers (vimentin). Western blot analysis was performed using antibodies against the indicated proteins. Samples were collected 10 days after induction of 16 candidate MET-TFs. Human mammary epithelial cells (MCF10A) were used as a positive control for epithelial lineages.

**Table 1.** List of candidate MET factors

| Symbol | Gene Name | Correlation with the E state |  |
| --- | --- | --- | --- |
|  |  | CCLE | FANTOM5 |
| *CDH1* | ankyrin repeat domain 22 | +++ | + |
| *EHF* | ETS homologous factor | +++ | – |
| *ELF3* | E74 Like ETS<br>Transcription Factor 3 | +++ | ++ |
| *FOXA1* | forkhead box A1 | +++ | +++ |
| *FOXA3* | forkhead box A3 | ++ | ++ |
| *GATA3* | GATA binding protein 3 | + | ++ |
| *GRHL1* | grainyhead-like 1 | +++ | +++ |
| *GRHL2* | grainyhead-like 2 | +++ | +++ |
| *GRHL3* | grainyhead-like 3 | +++ | +++ |
| *HNF1A* | HNF1 homeobox A | + | + |
| *IRF6* | interferon regulatory<br>factor 6 | +++ | ++ |
| *KLF4* | Kruppel-like factor 4 | + | + |
| *OVOL2* | ovo-like 2 | +++ | +++ |
| *TP63* | tumor protein p63 | + | ++ |
| *YBX2* | Y box binding protein 2 | ++ | +++ |
| *ZNF165* | zinc finger protein 165 | +++ | +++ |

“–” indicates no correlation; “+” indicates $r(\text{CDH1}) > 0$ and $r(\text{VIM}) < 0$ ; “++” indicates $r(\text{CDH1}) > 0.2$ and $r(\text{VIM}) < -0.2$ ; and “+++” indicates $r(\text{CDH1}) > 0.4$ and $r(\text{VIM}) < -0.4$ , respectively.

### Identification of OVOL2 as a potent MET inducer in dermal fibroblasts

The 16 selected TFs were then tested for their potential to induce MET in human neonatal dermal fibroblasts (HNDFs) by ectopic expression. When they were simultaneously introduced by pooled lentiviral vectors at a MOI of 20 (1.25 MOI for each TF), a subset of HNDFs started to form colonies with epithelial morphology (Figure 1B). The E marker proteins E-cadherin and cytokeratins were detected along with the morphological changes, whereas no changes were observed in the M marker vimentin (Figure 1C), likely because the epithelial conversion occurred in a small cell population, with most of the cells remaining in the mesenchymal state. These results suggest that the 16 candidate TFs contain factor(s) that can promote MET in fibroblasts.

We then looked for essential MET-TFs among the 16 candidates, by performing an “all-minus-one” screening based on Yamanaka’s approach, which was used when the minimum set of iPSC reprogramming factors was discovered by his group ^24^ (Figure 2A). This screening revealed that, among the 16 candidates, OVOL2 is the only indispensable factor for the induction of the E marker gene *CDH1*. However, OVOL2 alone was not sufficient to activate *CDH1* expression to the level achieved by all 16 factors (Figure 2B). Therefore, we hypothesized that other factors might activate *CDH1* expression cooperatively with OVOL2. To test this hypothesis, we performed an “OVOL2-plus-one” screening and found that HNF1A, TP63, and KLF4 exhibited the highest enhancement of OVOL2-induced *CDH1* expression (Figure 2C), which was confirmed to be statistically significant (Figure 2B).

**Figure 2.**
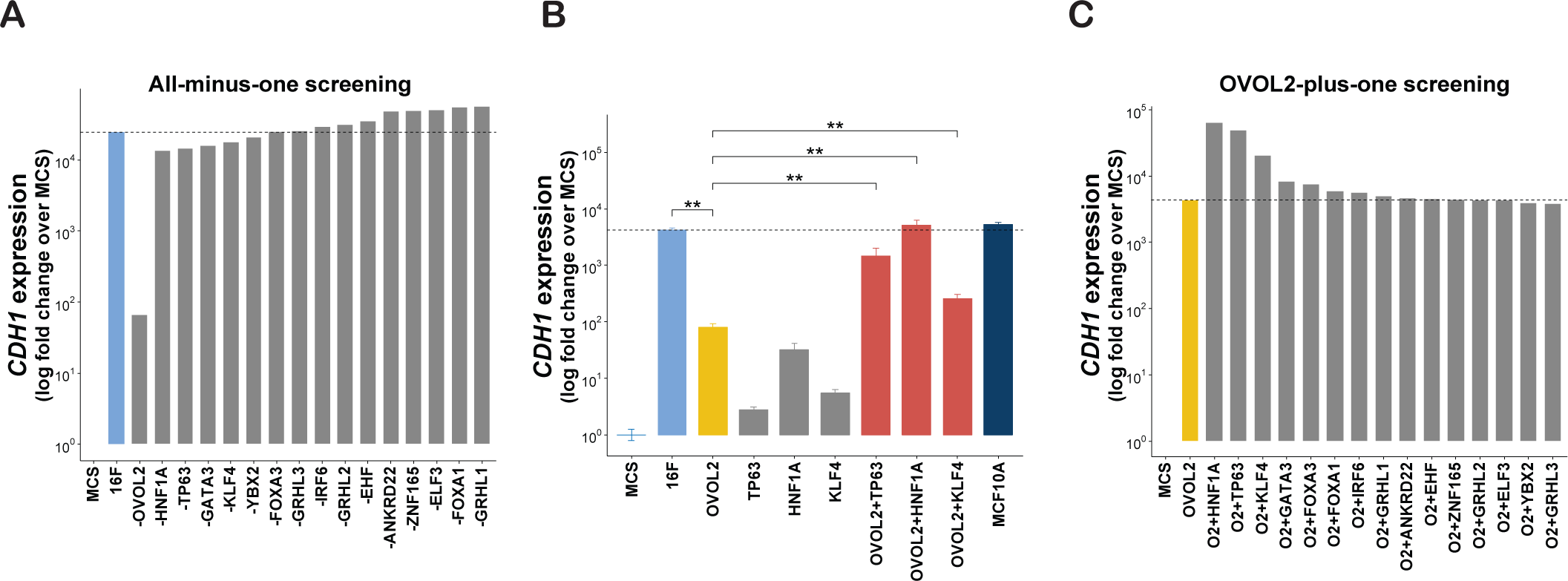
Identification of TF combinations that induce E phenotype in fibroblasts. (**A**) Identification of indispensable factor(s) for *CDH1* induction in HNDFs, using an all-minus-one screening approach. Quantitative RT-PCR (RT-qPCR) was performed to quantify *CDH1* expression 7 days after transduction of the indicated factors in HNDF. (**B**) Statistics analysis of *CDH1* induction by combination of OVOL2 with HNF1A, TP63 or KLF4. Results are presented as means ± S.D. of three biological replicates. \*\**p* < 0.01 (**C**) OVOL2-plus-one factor screening was performed to identify the cooperative partners of OVOL2 in inducing *CDH1* expression in HNDF.

### OVOL2 enhances reprogramming-mediated MET

The cooperative factors of OVOL2-induced *CDH1* expression, HNF1A, TP63, and KLF4, are involved in several processes of fibroblast reprogramming into epithelial lineages ^25–27^. Overexpression of TP63 together with KLF4 was reported to induce a keratinocyte-like state in human fibroblasts ^20^. In addition, previous studies used multiple TF combinations including HNF1A, HNF4A, and FOXA factors to reprogram human or mouse fibroblasts into hepatocyte-like state ^25,27^. Therefore, we hypothesized that reprogramming-mediated MET in keratinocyte- or hepatocyte-like states may be enhanced by OVOL2. To test this hypothesis, we investigated keratinocyte- and hepatocyte-like reprogramming using combinations of TP63/KLF4 (TK) and HNF1A/HNF4A/FOXA3 (HF), respectively, with or without OVOL2. A tetracycline-inducible lentiviral vector was generated using dual selection to control the expression of OVOL2 and TK or HF (See Methods section for details). To determine the cooperative enhancement of MET by the factor combinations, the TFs induction levels were set such that HF, TK, or OVOL2 could not, by themselves, show detectable morphological changes within 4–5 days after doxycycline (DOX) induction (bottom left or upper right in Figure 3A and B). However, under the same conditions, the combination of OVOL2 with either TK or HF (TK + OVOL2 or HF + OVOL2) dramatically facilitated epithelial-like morphological changes and became less proliferative (bottom right in Figure 3A and B). Further, both TK + OVOL2 and HF + OVOL2 combinations dramatically suppressed the directional migration phenotype, which is an important feature of MET ^28^, as compared with the respective controls including negative control vectors (Control), OVOL2 alone (OVOL2), TK alone (TK), and HF alone (HF) (Figure 3C). These data implicate that, in fibroblasts, functional MET is facilitated by the combination of TK + OVOL2 or HF + OVOL2.

**Figure 3.**
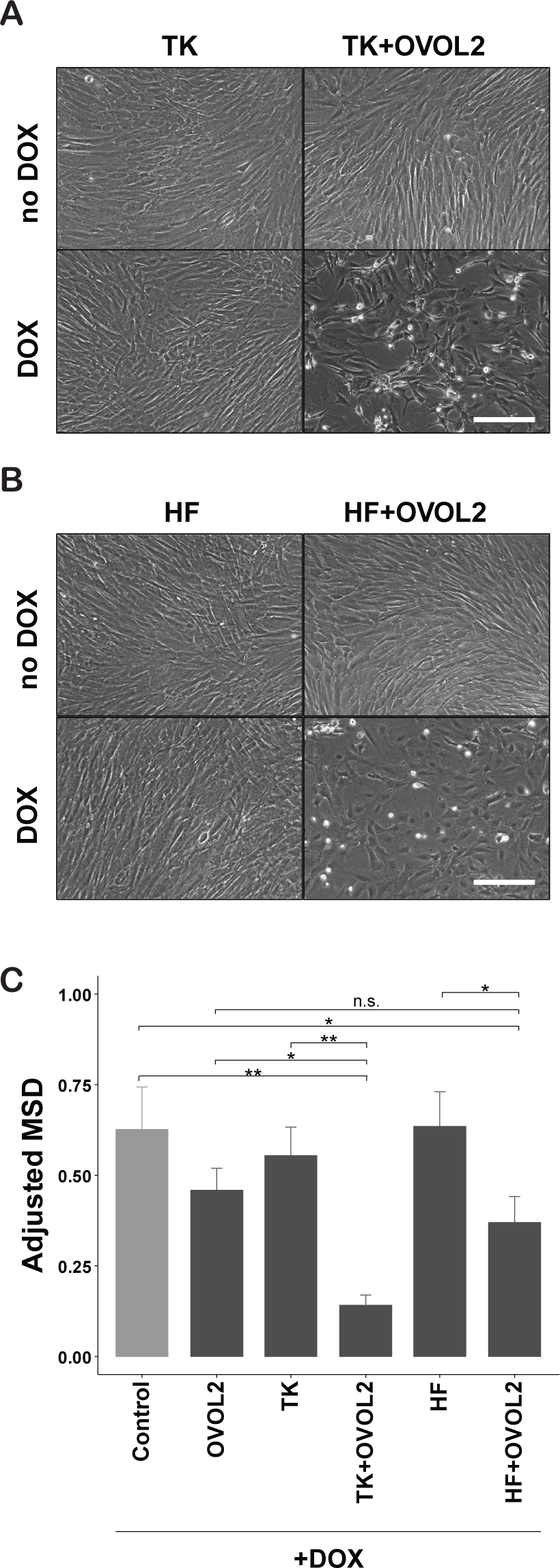
Cooperative induction of epithelial phenotype by combined OVOL2 and lineage factors. (**A–B**) Morphological changes induced by TFs combinations. Tetracycline-inducible (Tet-ON) puromycin-resistant lentivirus for KLF4/TP63 (TK) or HNF1A/HNF4A/FOXA3 (HF), Tet-ON hygromycin-resistant lentivirus for OVOL2, and the corresponding negative controls for each expression system were transduced in HNDF. After dual antibiotic selection of the infected cells, transduced genes were induced by doxycycline (DOX, 500 ng/mL) treatment for 4 days. Scale bar = 100 μm. (**C**) Motility phenotype changes induced by TFs combinations. The indicated TFs combinations were induced in HNDFs by DOX as described in (**A–B**). Four days after induction, DOX was removed and cellular movement and proliferation were tracked by nuclear RFP signals using the IncuCyte live-cell imaging system. The mean squared displacement (MSD) for the cellular movement was obtained as described in the Methods section.

### Cooperative induction of a keratinocyte-like state by the combination of OVOL2 and TP63/KLF4

Next, we examined whether OVOL2 facilitates fibroblast reprogramming into specific epithelial lineages. Since several previous studies reported the conserved role of the OVOL family factors during ectodermal tissue development ^29,30^, we investigated the involvement of OVOL2 in the reprogramming of HNDFs into the keratinocyte-like state. The results showed that TK + OVOL2-transduced HNDFs showed a dramatically increased expression of the keratinocyte-specific (K) marker genes *KRT14* and *TP63* (Figure 4A). Consistent with the changes in morphology/physiology (Figure 3), the E marker *CDH1* was dramatically induced by the combination of TK + OVOL2 (Figure 4A). Importantly, the expression of the M marker *VIM* was significantly reduced by TK + OVOL2, thus confirming M-state suppression concurrently to E-state activation (Figure 4A). Furthermore, transduction of TK + OVOL2 in HNDFs resulted in a dramatic increase of a KRT14 protein-expressing cell population at a 43% rate, whereas with TK, only 5.8% of cells became KRT14 positive (Figure 4B). This suggests that the reprogramming efficiency was increased at the population level. Taken together, these results support that the combination of TK + OVOL2 cooperatively facilitates HNDF reprogramming both at cellular and gene expression levels toward the keratinocyte-like state through activation of the keratinocyte-specific transcriptional regulatory network with MET enhancement.

**Figure 4.**
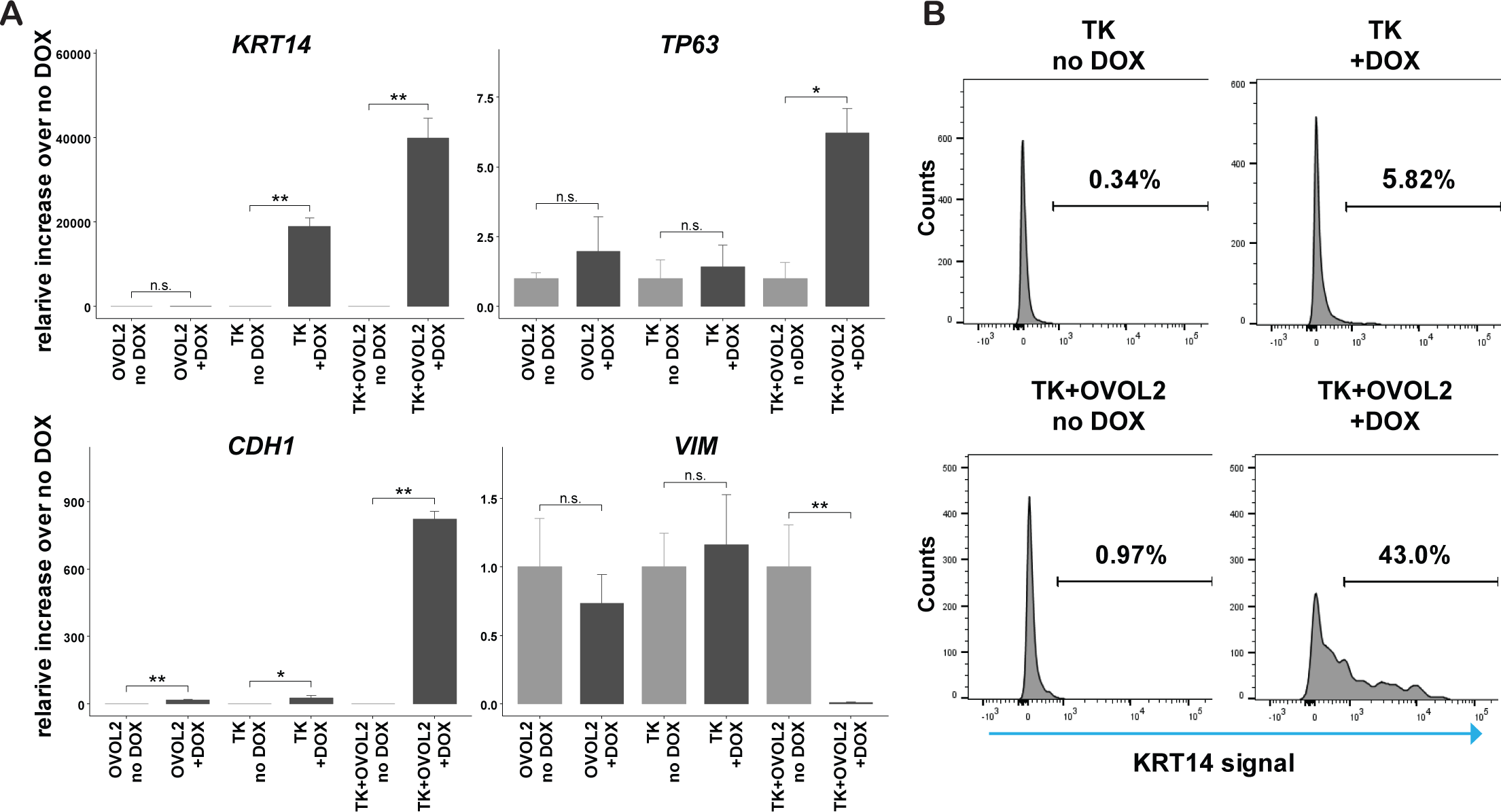
Enhanced induction of keratinocyte-like state by combinations of TP63/KLF4 and OVOL2. (**A**) Changes in marker gene expression upon inducible expression of TK and OVOL2. Keratinocyte lineage markers *KRT14* and *TP63*, an E marker *CDH1*, and an M marker *VIM* were quantified by RT-qPCR after DOX induction of the indicated factors in HNDF. (**B**) Induction of KRT14+ population upon inducible expression of TK and OVOL2. A keratinocyte marker KRT14-positive population was assessed by flow cytometry after DOX induction of the indicated factors in HNDF.

### The combination of TP63/KLF4 and OVOL2 enhanced transcription and chromatin accessibility changes toward the keratinocyte-like state

Chromatin state directly reflects the lineage identity of a given cell, providing clues to understand the regulatory mechanisms of cell-state transitions ^31^. Therefore, we evaluated changes in chromatin accessibility following induction of TF combinations in HNDFs, using ATAC-seq ^32^. Cooperative induction of open or closed chromatin state by TK + OVOL2 was observed at the promoter region of the representative E or M gene loci, such as *CDH1* or *VIM*, respectively (Figure 5A, marked in red). Interestingly, roadmap-defined *CDH1* enhancers ^33^ showed an open chromatin state only under OVOL2 expression, whereas *VIM* enhancers showed a tendency toward the closed chromatin state by the combination of TK + OVOL2 (Figure 5A, marked in blue). This suggests that those loci are involved in the cell reprogramming mechanism. We then compared chromatin state changes with transcriptomic changes assessed by CAGE ^34^. At a genome-wide scale, TK + OVOL2-transduced HNDFs showed greater changes toward keratinocyte-like state than either OVOL2 or TK alone, both in CAGE and in ATAC-seq data (Figure S3). Next, we created a list of E, K, M, fibroblast (F)-specific gene sets based on previous literature (Table S1) ^35–38^ and investigated how these gene categories would behave transcriptionally and how their corresponding chromatin accessibility would behave epigenetically during reprogramming (Figure 5B). The combination of TK + OVOL2 markedly enhanced both the transcriptional activity and the chromatin accessibility of E and K genes. In contrast, both OVOL2 and TK alone failed to induce expression and chromatin accessibility in E and K genes, apart from a mild activation of E gene expression by OVOL2 alone. On the other hand, the expression of M genes was inhibited by either OVOL2 or TK alone, which was further enhanced by the combination of TK + OVOL2, whereas chromatin accessibility changes were observed only for TK + OVOL2. Transcriptional inhibition of F genes was observed only in the TK + OVOL2-introduced HNDFs. However, chromatin accessibility did not show significant changes. These results indicate that TK + OVOL2 substantially enhances epigenetic changes of HNDFs toward keratinocyte-like state at representative genomic loci, and most of transcriptional changes are accompanied by changes in the chromatin state.

**Figure 5.**
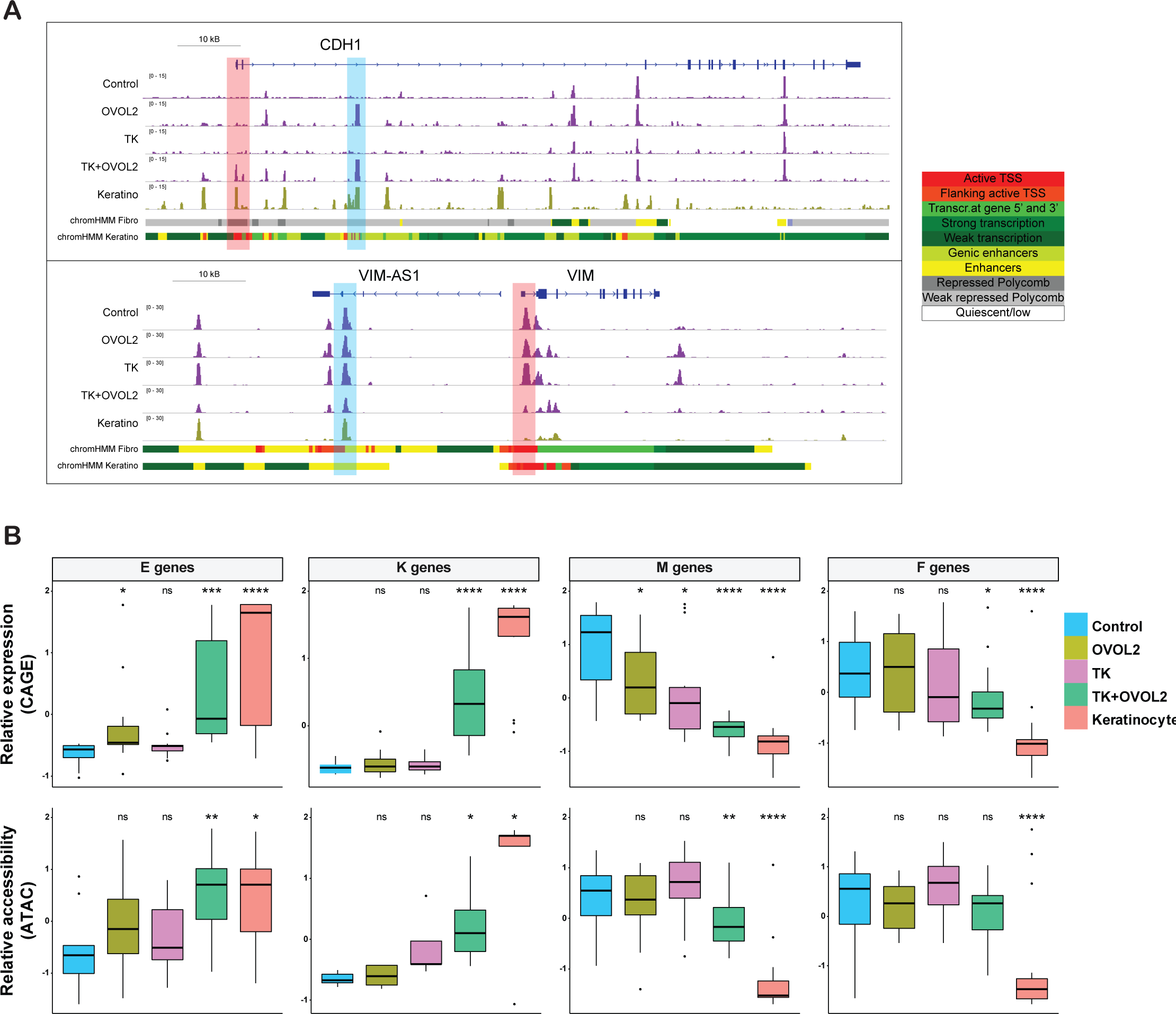
Enhanced chromatin accessibility changes by OVOL2 combined with lineage factors. (**A**) Genome browser view of ATAC-seq signals at the *CDH1* and *VIM* loci. Promoter and potential enhancer regions that showed enhanced changes by a combination of TK + OVOL2 are highlighted in red and blue, respectively. Roadmap annotation of genomic regions (promoters and enhancers for the indicated cell types) is shown at the bottom. (**B**) A box plot showing Z-scores for expression (CAGE, top) or chromatin accessibility (ATAC, bottom) in E, M, K, and F genes, respectively. *p*-values were calculated through comparison with control samples. \**p* < 0.01, \*\**p* < 0.001, \*\*\**p* < 0.0001, \*\*\*\**p* < 0.00001.

### Distinct regulation of chromatin accessibility by OVOL2 and TP63/KLF4

Chromatin accessibility evaluation is a powerful approach to dissect the activity of regulatory elements through TF-binding motif analysis ^39^. To understand the transcriptional regulatory networks that define fibroblast and keratinocyte lineages, we investigated the differences in enriched TF-binding motifs at open chromatin loci between primary fibroblasts (i.e., control HNDF) and primary keratinocytes using the analytical program chromVAR ^40^. Among the 808 curated TF motifs used in the analysis, 34 were significantly more accessible in fibroblasts whereas 72 were significantly more accessible in keratinocytes (q < 0.01 and enrichment_score > 5) (Figure 6A, Table S2). Among them, 9 out of 34 and 18 out of 72 showed a significantly different expression in the CAGE data (q value <0.01 and |log fold change| > 1; Figure 6B). This unbiased analysis revealed that TP63 and KLF4 motifs were significantly more accessible in keratinocytes, indicating the importance of these factors to specify the keratinocyte lineage (Figure 6A). In contrast, OVOL1/2 motifs were significantly more accessible in primary fibroblasts (Figure 6A). When directly compared with the CAGE expression data (Figure 6C), the gene expression and motif enrichment scores of OVOL1/2 show opposite trends, in that OVOL1/2 motifs become inaccessible when OVOLs are expressed, suggesting that OVOLs act as negative regulators of the chromatin state. Other factors of interest included the AP1 family factors JUN/JUND/FOS, inhibitor of DNA binding family ID3/4, and EMT-TFs SNAI1 and ZEB1. Notably, no other candidate MET-TFs presented in Table 1 showed significant differences between the two cell lineages regarding their motif accessibilities (Table S2).

**Figure 6.**
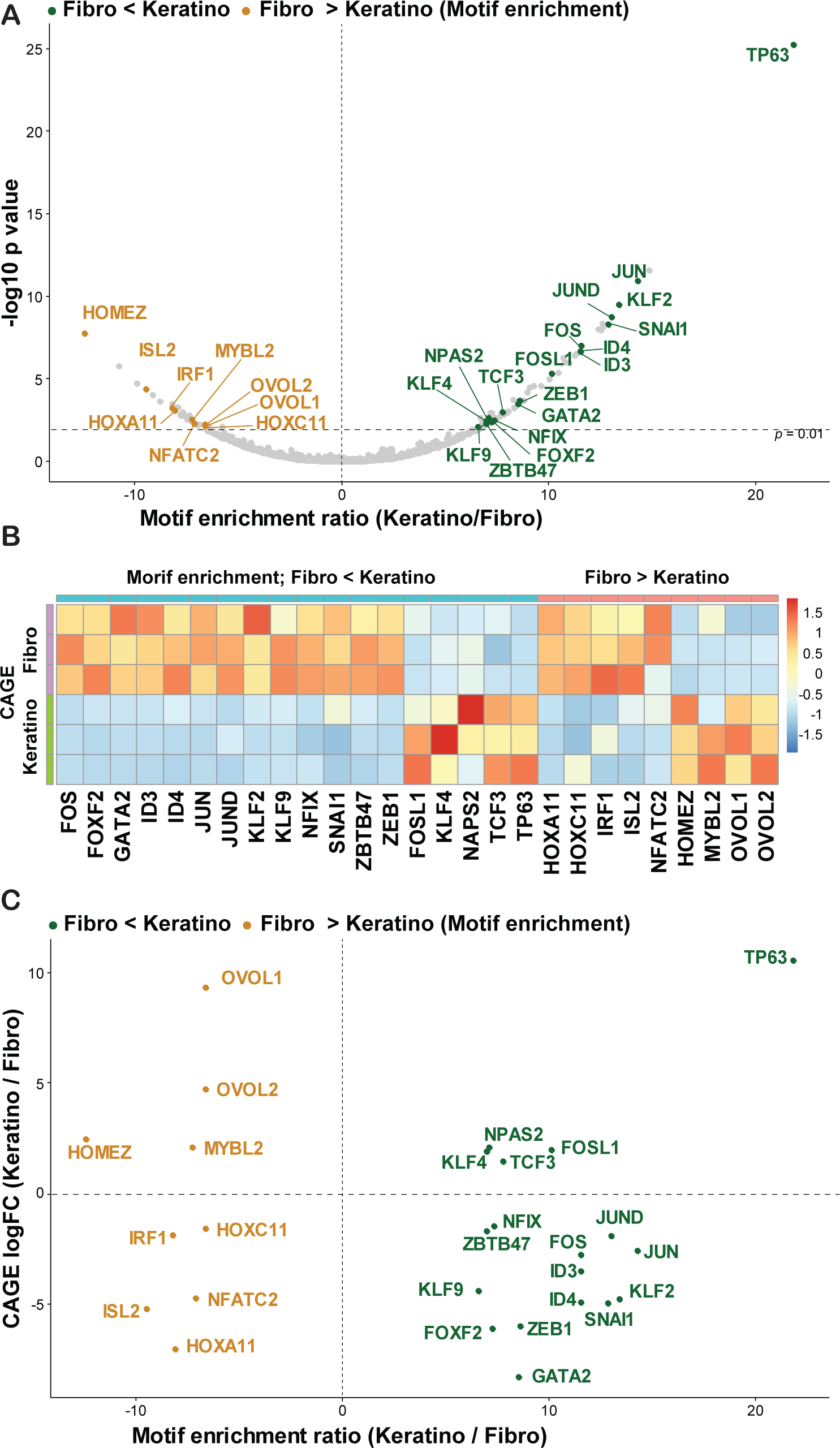
Motif accessibility and TFs expression between fibroblast- and keratinocyte-lineages. (**A**) Top differentially accessible TF motifs between fibroblasts and keratinocytes. TFs that showed significant differences in CAGE expression (*q* < 0.01 and |log fold change| > 1) were labeled. (**B**) Heatmap for the expression levels of the indicated TFs in FANTOM5 CAGE data. Three replicates for primary dermal fibroblasts and keratinocytes are shown. (**C**) A scatter plot of TFs by motif enrichment score (*=Z* score, *x* axis) and CAGE expression fold changes (*y* axis).

Next, we investigated how these factors change their motif accessibilities during reprogramming. In a t-SNE plot generated by the motif enrichment scores, TK + OVOL2-treated cells became closer to primary keratinocytes than control, OVOL2, or TK-treated cells (Figure 7A). We then compared the motif enrichment of individual factors in open chromatin peaks for each condition (Figure 7B). The OVOL2 motif became inaccessible in TK + OVOL2-treated HNDFs at a level similar to keratinocytes, and its expression level increased in TK + OVOL2-treated HNDFs. TP63 expression and motif enrichment was dramatically elevated in keratinocytes and elevated to a lesser extent in TK + OVOL2-treated HNDFs. Overall, KLF4 showed a consistent increase in motif enrichment and gene expression during reprogramming toward keratinocyte-like state, except for an increase in the transcript observed in TK alone, which could have been caused by contamination of exogenously introduced sequences. Overall, a positive correlation was observed between the expression and motif enrichment of TP63 and KLF4. In addition to reprogramming TFs, ZEB1 and ID3 showed similar reciprocity between gene expression and motif activity, as these factors were highly expressed in control HNDFs, but motifs were enriched in activated peaks of TK-OVOL2-treated HNDFs and keratinocytes (Figure S4). These results indicate that OVOL2 and TK regulates the MET process through distinct chromatin regulation mechanisms (negative vs positive), which may have caused enhanced transition of fibroblasts into epithelial lineages.

**Figure 7.**
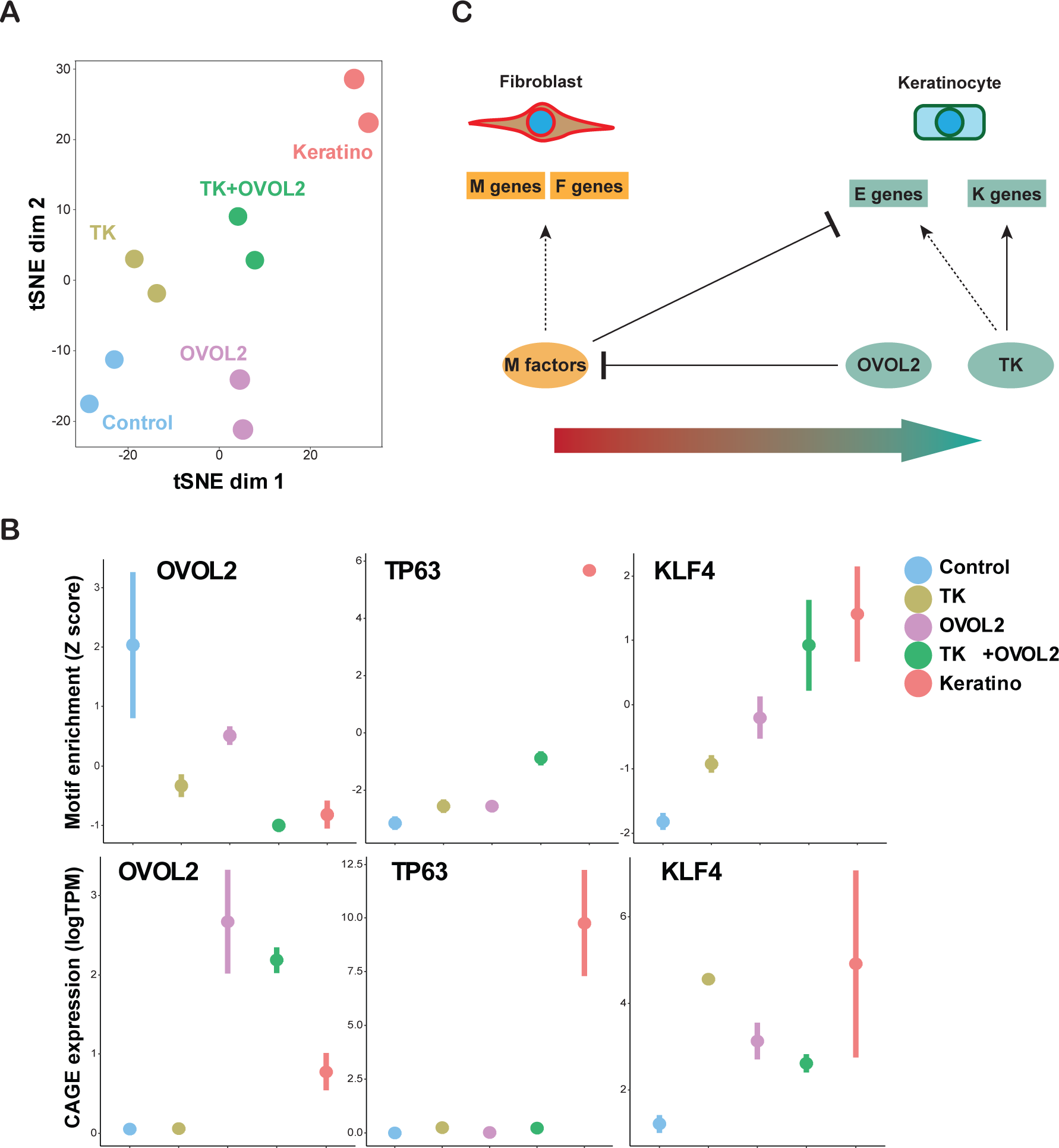
Changes in motif accessibility and TF expression during reprogramming. (**A**) tSNE plot using TF motif enrichment scores for each condition based on the ATAC-seq data. (**B**) Motif accessibility and TF genes expression. Motif enrichment score (Z score) and CAGE expression (logTPM) are shown in the top and bottom panels, respectively, for the indicated TFs. (**C**) A proposed model for MET induction in fibroblast by OVOL2 and lineage factors.

## DISCUSSION

### Identification of OVOL2 as a potent MET inducer in dermal fibroblasts

The present study aimed to identify potent MET-TFs in fibroblasts. Our candidate list contained factors previously reported as regulators of MET, the GRHL family factors ^11,12^ ELF3 ^16^, GATA3 ^18^, FOXA1/3 ^19^, KLF4 ^20^, and OVOL2 ^13–15^. However, among them OVOL2 was the only essential factor to induce E genes in our screening (Figure 2). OVOL2 and its homologue OVOL1 (not included in this study due to a technical reason) were previously reported to induce MET in various types of cancer cells ^13^ and to protect the epithelial identity of normal epithelial cells ^14,15^. The current study extends these findings to fibroblasts, a non-epithelial cell type with an established M state. The fact that other candidate TFs with potential MET-inducing activities in other cell types did not show any potent effect in fibroblasts may implicate that epithelial programs are strongly suppressed in fibroblasts where OVOL2 can only break this barrier. Consistently, OVOL2 was the only factor showing differential motif accessibility in ATAC-seq analysis, among the candidate factors (Table S2).

Mechanistically, the ATAC-seq results indicated that OVOL2 induces a closed chromatin state at its target loci, which is consistent with a previous report demonstrating that OVOL2 acts primarily as a transcription repressor in epithelial cells ^15^. This indicates that OVOL2 activates E (and K) genes indirectly. How does OVOL2 activate these genes? ATAC-seq analysis revealed that TFs’ motif-containing regions, including ZEB1 and ID3, are significantly accessible in keratinocytes and become accessible in fibroblasts upon introduction of TK + OVOL2. ZEB1 is an EMT-TF that plays an essential role for the maintenance of the M phenotype. In fact, previous studies have demonstrated that ZEB1 is a repressor of many E genes including *CDH1* ^41^ and constitutes an important target of *OVOL2* ^15^. On the other hand, ID3 exhibits a broad biological effect and its involvement in MET has not been clearly demonstrated ^42^. Therefore, a potential mechanism underlying OVOL2-mediated E gene activation resides in inducing suppression of the ZEB1 repressor (= de-repression of E genes), which may comprise a prerequisite of MET induction in fibroblasts.

### Reprogramming-mediated MET was enhanced by OVOL2

Our screening also identified OVOL2 as a cooperative MET inducer, when introduced together with the epithelial reprogramming factors HNF1A, TP63, and KLF4. Our results indicate that TP63/KLF4 directly activate their targets, including E and K factors at a chromatin level, as their expression and motif accessibility were positively correlated (Figures 6 and 7). Therefore, TP63/KLF4-mediated MET must occur via mechanisms different from OVOL2. These factors therefore use independent pathways to activate E genes, resulting in a cooperative MET induction in fibroblasts.

The combined effect of OVOL2 and TP63 may be relevant in physiological settings, since their expression patterns are important for the development and maintenance of ectodermal tissues. For example, OVOL2 and TP63 are co-expressed in the basal layer of the interfollicular epidermis ^14^, but mutually exclusively expressed in mammary epithelium, since OVOL2 is preferentially expressed in luminal layers ^15^, whereas TP63 is specifically expressed in basal myoepithelial cells ^43^. This might be related with the fact that mammary basal cells (i.e., myoepithelial cells) exhibit less epithelial phenotypes than basal keratinocytes. Further *in vivo* studies are needed to investigate how these factors interact to regulate epithelial fate.

### OVOL2 facilitates cell-state changes of fibroblasts towards specific epithelial lineages

Our results also indicate that OVOL2-mediated MET facilitates cell-state changes toward specific epithelial lineages such as keratinocytes. In this context, our previous reports demonstrated that suppression of the fibroblast-specific TF network facilitates the direct reprogramming of fibroblasts into specific lineages such as adipocytes or monocytes ^38,44^. This suggests that, in direct cell reprogramming, suppression of original cell-state is critical to activate the TF network of the target state. Along this line, it has been reported that miR200 families also enhance iPSC induction through EMT-TFs suppression in fibroblasts ^3^. Similarly, our results indicate that exogenous OVOL2 expression suppresses the original M state in fibroblasts, promoting the transition toward the E state (Figure 7C). Therefore, suppression of the original M state may be a key event to enhance the epithelial reprogramming of fibroblasts.

### Conclusions

The present study revealed the potent MET-inducing role of OVOL2 in fibroblasts and showed that TP63/KLF4-mediated MET was further enhanced by OVOL2. In addition, we demonstrated that OVOL2-mediated MET facilitates the TP63/KLF4-mediated direct cell reprogramming of fibroblasts towards the keratinocyte-like state. Finally, transcriptome/epigenome analysis indicated that OVOL2 and TP63/KLF4 function differentially at the epigenetic level. Since MET is crucial during TF-induced cell reprogramming of fibroblasts into epithelial lineages, further understanding of MET induction mechanisms holds great potential to improve cell reprogramming technologies for future clinical applications.

## METHODS

### Cell culture and transdifferentiation

HNDF (Lonza, Switzerland) were grown in αMEM (Thermo Fisher, USA) supplemented with 10% fetal bovine serum. Primary keratinocytes (Lonza) were grown in KGM-Gold™ Keratinocyte Growth Medium (Lonza). For TF-mediated reprogramming into keratinocyte-like state, culture media were switched from αMEM to KGM-Gold™ after introduction of the indicated TFs. For TF-mediated reprogramming into hepatocyte-like state, culture media were switched from αMEM to HGM™ Hepatocyte Growth Medium (Lonza) and cells were plated onto collagen I-coated plates (Thermo Fisher) at the first passage after TF induction.

### Lentiviral expression systems

Candidate MET-TFs with cDNAs available in our cDNA collection were selected (Table 1). The initial screening was performed by cloning cDNAs into CSII-EF-RfA-IRES2-Puro lentiviral construct ^45^. Inducible expression of TP63, KLF4, HNF1A, HNF4A, and FOXA3 was achieved using a lentiviral expression construct with puromycin resistance (pCW57.1, a gift from David Root, Addgene plasmid # 41393). Inducible expression of OVOL2 was achieved using a construct with hygromycin resistance (pSLIK-Hygro, a gift from Iain Fraser, Addgene plasmid # 25737). Optimal transduction efficiency with low multiplicity of infection (MOI) was achieved by dual selection using hygromycin- and puromycin-resistant constructs. Lentiviruses were prepared in a mixture of the following packaging constructs: pCMV-VSV-G-RSV-Rev and pCAG-HIVgp (provided by RIKEN BRC, Japan). The nuclear red fluorescent protein (RFP)-expressing lentiviral construct (LV-RFP) was a gift from Elaine Fuchs (Addgene plasmid # 26001). To infect lentiviruses, HNDFs were seeded in a 6-well plate at a density of 1 × 10^6^ cells/well, followed by transduction, performed on the second day by adding 8 µg/mL polybrene and 10 MOI of lentivirus into each well. The antibiotics selection was started 2 days after transduction.

### Immunocytochemistry

Cells grown in 8-well-chambered slide (Thermo Fisher) were fixed in 4% paraformaldehyde, then permeabilized with 0.2% Triton X-100. After washing and blocking, cells were stained with rabbit polyclonal anti-β-catenin antibody (1:1000, H-102, Santa Cruz, USA) and labeled with the secondary antibody conjugated with Alexa-Fluor-488. Counterstaining was performed with DAPI.

### Western blotting

Total protein was extracted from cells by lysing with radioimmunoprecipitation assay buffer (Takara, Japan). Ten micrograms of protein were run on 4–20% gradient polyacrylamide gels (Thermo Fisher) and transferred to nitrocellulose membranes (Merck, Germany). The membranes were then blocked with 5% nonfat milk in Tris-buffered saline with Tween20 (10 mM Tris, pH 8.0, 150 mM NaCl, 0.5% Tween20) and probed with the following antibodies: anti-E-cadherin (BD Biosciences, USA, 1:1000), anti-pan-cytokeratin (Wako, Japan, 1:1000), anti-Vimentin (Cell Signaling Technologies, USA, 1:1000), and anti-β-actin (Sigma-Aldrich, USA, 1:5000).

### Cell physiology assays

Cell motility was quantified by image-based analysis using a live-cell imaging device, the IncuCyte (Essen Bioscience, USA). Cellular movement was tracked by nuclear RFP signals from HNDFs that had been infected with LV-RFP. Cell images were taken every 15 min, and each fluorescent spot was tracked using the Fiji Trackmate. Each spot’s coordinates at each time point were used to calculate the mean squared displacement (MSD) using the following formula: 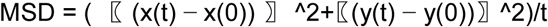. Within each track, the values for all time points were filtered and the maximum MSD was kept and divided by the time duration of the track to adjust for time dependency. The time-adjusted MSD values for all tracks were averaged to obtain the representative MSD value for each cell group.

### Quantitative RT-PCR (RT-qPCR)

Total RNA was isolated using a QIAGEN miRNeasy mini kit (QIAGEN, Germany), and first-stranded cDNA was prepared using the Prime Script RT kit (Takara) according to the manufacturer’s instructions. Real-time PCR was performed using a StepOnePlus™ real-time PCR system (Thermo Fisher), with SYBR Premix Ex Taq™ II (Takara). Comparative analysis was performed for the genes of interest, which were normalized by the housekeeping genes GAPDH and ACTB. The primer sequences used in this study are presented in Table S3.

### Cap analysis of gene expression (CAGE)

CAGE libraries were prepared as previously described ^46^. Briefly, 3 μg of total RNA from each sample were subjected to reverse transcription, using SuperScript III Reverse Transcriptase (Thermo Fisher) with random primers. The 5-end cap structure was biotinylated by sequential oxidation with NaIO_4_ and biotinylation with biotin hydrazide (Vector Laboratories, USA). After RNase I treatment (Promega, USA), the biotinylated cap structure was captured with streptavidin-coated magnetic beads (Thermo Fisher). After ligation of 5’ and 3’ adaptors, second-strand cDNA was synthesized with DeepVent (exo-) DNA polymerase (New England BioLabs, USA). The double-stranded cDNA was treated with exonuclease I (New England BioLabs) and purified. The resulting CAGE libraries were sequenced using single-end reads of 50 bp on the Illumina HiSeq 2500 (Illumina, USA). CAGE data for the primary keratinocytes were obtained from the FANTOM5 data resource (http://fantom.gsc.riken.jp/5/sstar/Main_Page). The extracted CAGE tags were then mapped to the human genome (hg38). After filtering low-quality reads, mapped CAGE tags were counted regarding FANTOM5-CAT TSSs ^47^, providing a unit of CAGE tag start site. The tags per million (tpm) were calculated for each TSS peak and computed as gene expression levels for multiple TSS peaks associated with a single gene.

### ATAC-seq

ATAC-seq was performed as previously described, with some modifications ^32^. Samples stored in the Cell banker (Takara) at −80 °C were used in this study. After thawing, the cell number and viability were quantified with Countes (Thermo Fisher), and the nuclei were extracted from approximately 25,000 live cells with CSK buffer (10 mM PIPES pH 6.8, 100 mM NaCl, 300 mM sucrose, 3 mM MgCl_2_, 0.1% Triton X-100) on ice for 5 min. Transposase reaction was then performed, followed by 9 to 12 total cycles of PCR amplification. Amplified DNA fragments were then purified with the QIAGEN MinElute PCR Purification Kit (QIAGEN) and size-selected twice with Agencourt AMPure XP (1:1.5 and 1:0.5 sample to beads; Beckman Coulter, USA). Libraries were quantified by KAPA Library Quantification Kit for Illumina Sequencing Platforms (KAPA Biosystems, USA), and size distribution was determined by Bioanalyzer (Agilent Technologies, USA). The resulting ATAC-seq libraries were sequenced using single-end reads of 50 bp on the Illumina HiSeq 2500 (Illumina).

Kundajelab pipeline for ATAC-seq (https://github.com/kundajelab/atac_dnase_pipelines) was used for read processing, peak calling, and quality control ^33^. Briefly, fastq files were grouped together and input separately into the pipeline. After mapping to hg38 reference genome, peaks were called by MACS2 ^48^ for both individual samples and pooled replicates. A reference peak file was created by merging all pooled peaks from all four conditions and removing gaps smaller than 100 bp. After merging all detected ATAC-seq peaks in all cell types and conditions, a total of 254,578 open chromatin regions were detected. Finally, the raw reads in the reference peak regions were counted individually for each sample, using the multicov function of the bedtools software v2.27.0 (https://bedtools.readthedocs.io/en/latest/). For gene-based analysis, each peak was assigned to the closest gene.

### Hierarchical clustering and motif analysis of CAGE genes and ATAC peaks

Hierarchical clustering analysis was performed using the average linkage algorithm with a Spearman correlation distance matrix. Motif enrichment analysis was performed using an R package chromVar using the default set of motifs with some modifications ^40^. *Z* score for significance of motif enrichment was used as enrichment score.

### Statistics

To evaluate the significance of differences in gene expression (RT-qPCR/CAGE), motility phenotype, and chromatin accessibility (ATAC-seq), a two-tailed Student’s *t*-test was used to calculate *p*-values.

## ACKNOWLEDGEMENTS

We thank Chung-Chau Hon for useful advice in data analysis and Mona Tang for technical assistance. This work was supported by Takeda Science Foundation, Grant-in-Aid for Scientific Research (KAKENHI) on Innovative Areas “Cellular Diversity” (JP18H05106) and KAKENHI (JP16KK0165) to KW. This work was also supported by a research grant from the Ministry of Education, Culture, Sport, Science and Technology of Japan for the RIKEN Center for Integrative Medical Sciences.

## AUTHOR CONTRIBUTIONS

KW, AM, and HS designed the research. KW, YL, MM, MK, and HN performed the experiments. KW, SN, JC, and KH analyzed the data. KW and HS wrote the paper. All authors approved the final version of this manuscript.

## DATA AVAILABILITY

Data for CAGE and ATAC-seq have been deposited with GEO under the access number GSE122036.

## SUPPLEMENTARY INFORMATION

Table S1 List of E-, K-, M-, and F-specific marker genes

Table S2 Accessibility score of TF motifs

Table S3 Primers used in this study

Supplementary Figures

