## Supplementary text for "OVOL2 induces mesenchymal-to-epithelial transition in fibroblasts and enhances reprogramming to epithelial lineages"

**Supplemental tables**

**Table S3. Primers used in this study**

| **Target gene** | Forward primer |  | Reverse primer |
| --- | --- | --- | --- |
| *CDH1* | AAAGGCCCATTTCCTAAAAACCT |  | TGCGTTCTCTATCCAGAGGCT |
| *VIM* | GACGCCATCAACACCGAGTT |  | CTTTGTCGTTGGTTAGCTGGT |
| *KRT14* | GCGGCCTGTCTGTCTCAT |  | TGAGCCGCATTCTGAACGAG |
| *TP63(endogenous)* | GGTCCCCACAGAGCAAGA |  | TGCAATGACAGCCCTTGA |

**Supplemental figure legends**

**Figure S1. Correlation between the expression of representative E and M markers.**

(**A**) Analysis with CCLE datasets. Pearson correlation coefficient r was calculated between each transcript and representative E genes (CDH1, EPCAM, KRT19, and TJP1) and M genes (VIM, MMP3, CDH2, and FN1) across 1038 microarrays for cancer cell lines. The correlation coefficient r for each transcript between the indicated combinations of E and M markers is shown in the scatter plots. (**B**) Analysis with FANTOM5 datasets. Expression correlation was analyzed across 1829 CAGE data for normal cells, as described in (**A**).

**Figure S2. Expression correlation analysis of FANTOM5-defined TFs with FANTOM5 CAGE datasets.**

Among 1995 FANTOM5-defined TFs, 1675 TF transcripts were identified in the FANTOM5 datasets. The Pearson correlation coefficient *r* was calculated between each transcript and *CDH1* or *VIM* across 1829 FANTOM5 CAGE data for normal cell types. Representative EMT-TFs (blue) and 16 candidate MET-TFs (red) were labeled.

**Figure S3. Heatmap and unsupervised clustering by CAGE and ATAC-seq.**

Heatmap with unsupervised clustering of the indicated samples and annotated marker genes for E, M states and keratinocytes by CAGE (**A**) and ATAC-seq (**B**). The top differentially expressed genes and differentially regulated ATAC-seq peaks between control fibroblasts (Control) and primary keratinocytes were used for the analysis. Note that all replicate pairs are co-aggregated at the first level of the dendrogram and the TK + OVOL2 samples changed their expression pattern toward keratinocytes.

**Figure S4. Changes in motif accessibility and expression of TFs during reprogramming (ZEB1 and ID3).**

Motif accessibility and TF genes expression. Motif enrichment score (*Z* score) and CAGE expression (logTPM) are shown in the top and bottom panels, respectively, for the indicated TFs.
