## Supplementary figures and images for "OVOL2 induces mesenchymal-to-epithelial transition in fibroblasts and enhances reprogramming to epithelial lineages"

### Supplementary figures

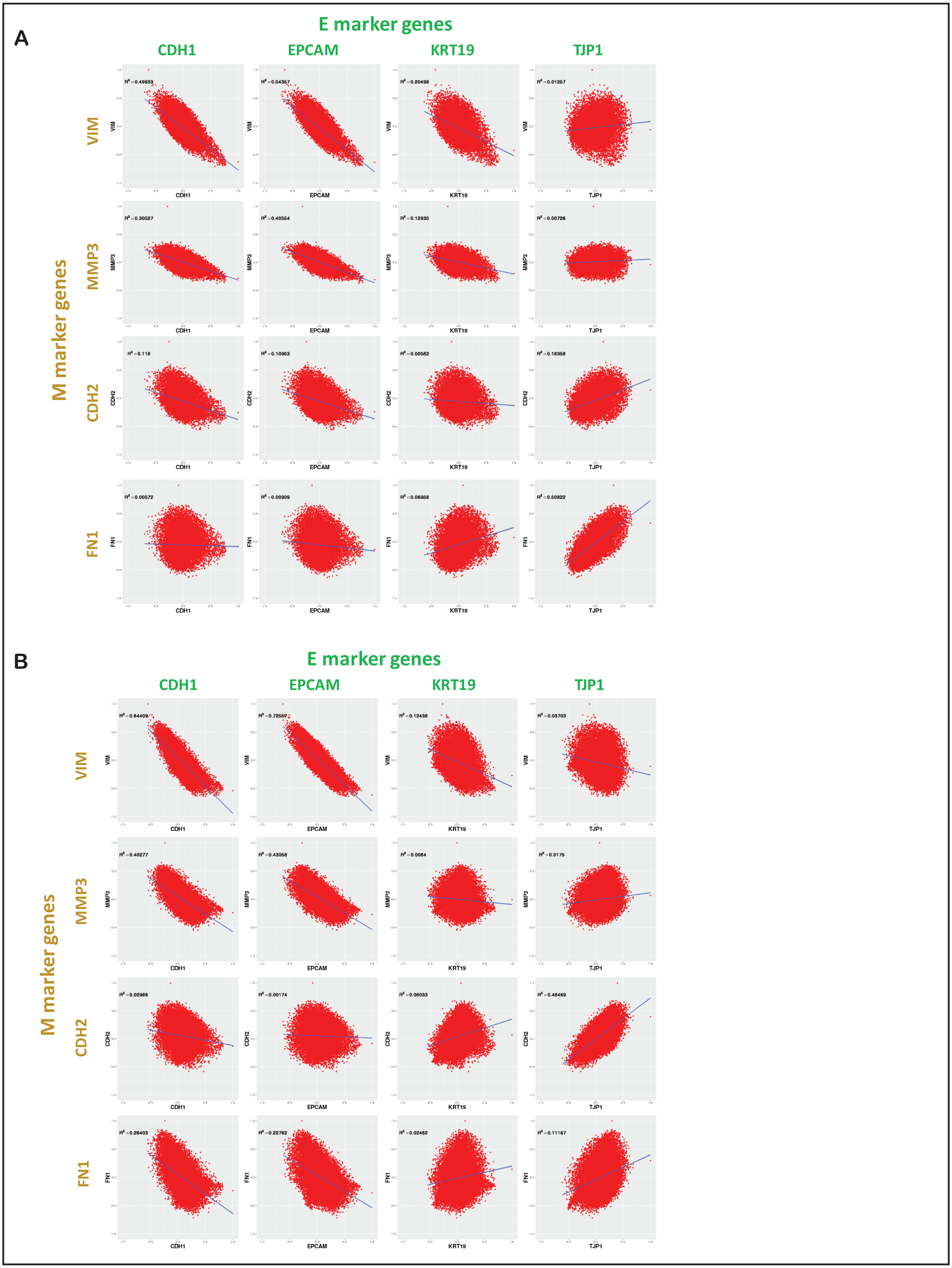

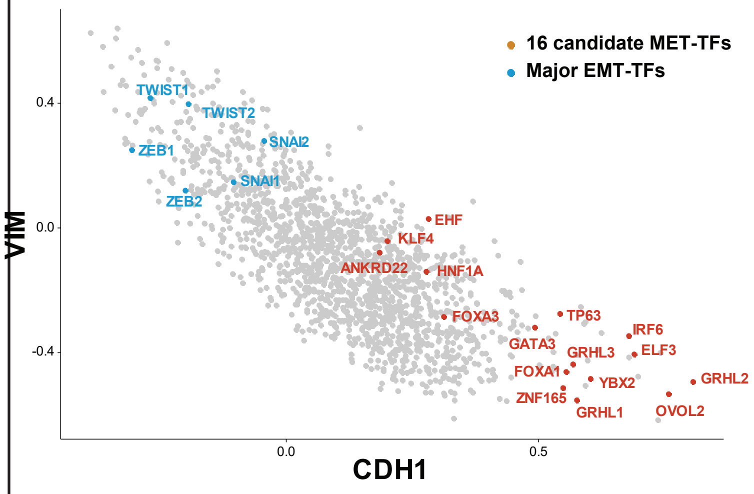

#### Figure S3

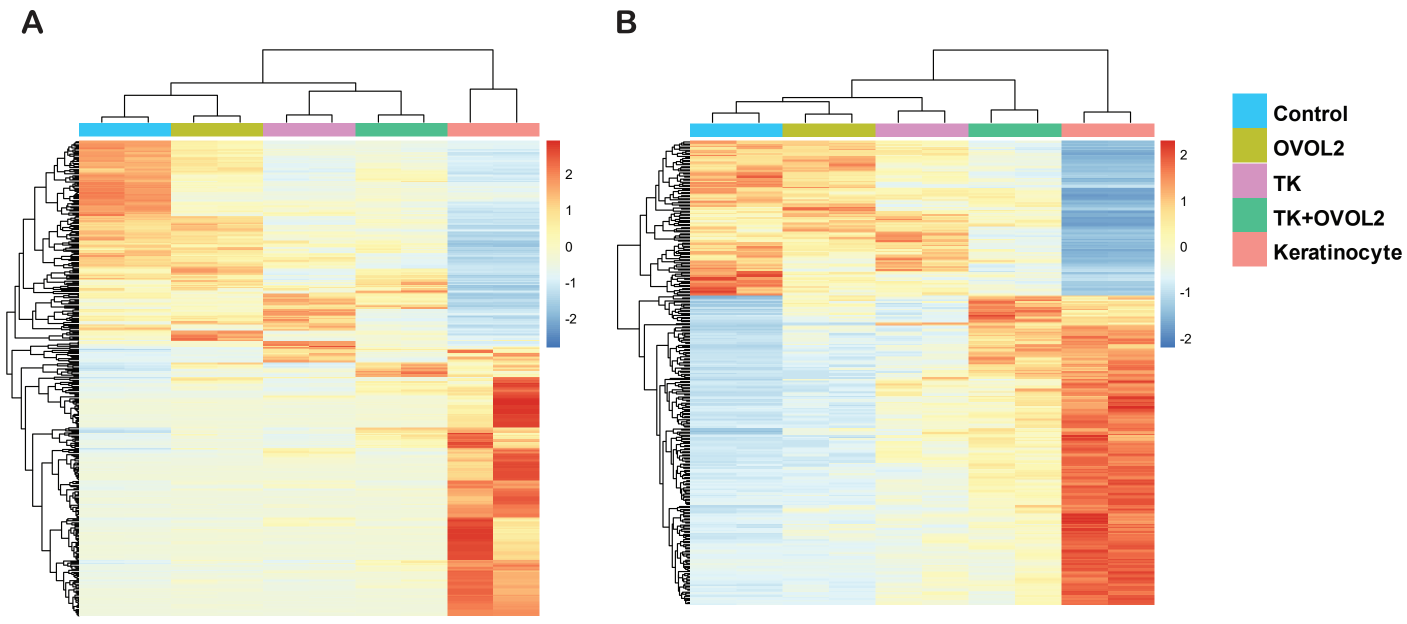

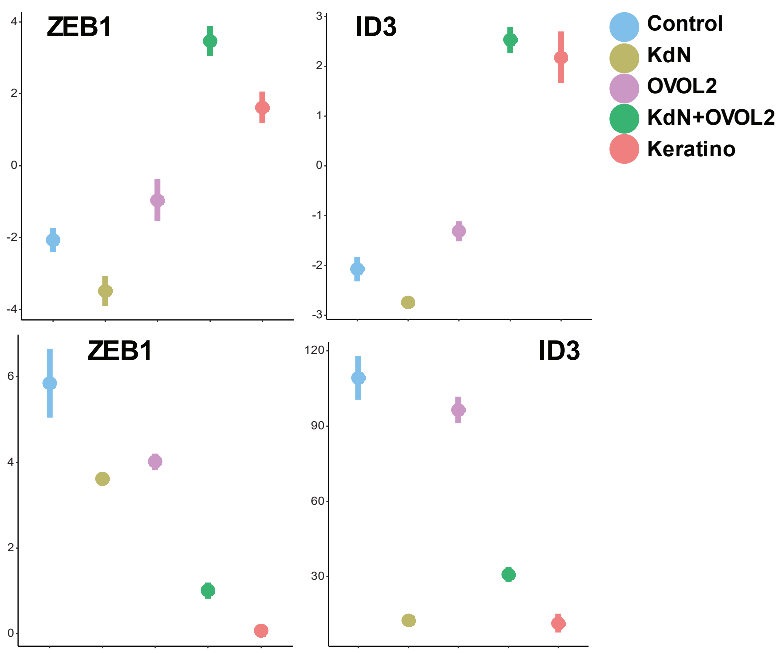
